# A tale of three morphs I: Variations in nectar, reproductive compatibility, and abundance explain reproductive success of the morphs in a polymorphic ginger from Western Ghats, India

**DOI:** 10.1101/2024.09.21.614269

**Authors:** Saket Shrotri, Vinita Gowda

## Abstract

**Background and aims:** Floral color polymorphism is common in angiosperms, yet in perennial, clonal plants, proximate trait differences and reproductive differences among sympatric morphs remain poorly understood. This is primarily because perennial individuals can carry signatures of past selection regimes. We studied bract color polymorphism in a nocturnal ginger *Curcuma caulina* to ask: (i) Do nectar rewards, reproductive compatibility and pollinator visitation differ among color morphs? (ii) Does morph abundance correlate with fitness via ecological (pollinator-mediated) or physiological (compatibility) pathways? (iii) Do these traits predict realized reproductive success?

**Methods:** We measured 22 floral/vegetative traits (n=30 per morph), nectar rewards (106 flowers), pollinator visitation rates (648 hours), and physiological compatibility (471 hand-pollinations treatments) and natural fruit and seed set (≥50 per morph) across three dominant morphs. Path analysis tested direct and indirect effects of morph abundance, nectar traits, pollinator visitation, and compatibility on reproductive fitness.

**Key results:** Morphs did not differ in their morphological traits but showed differential nectar and reproductive compatibility traits. The rare red-white morph produced the highest nectar energy, received the most visits, had a leaky self-compatibility, and yet showed lowest seed count per fruit, while the most common green-red morph was self-incompatible, showed higher cross-compatibility and had highest seeds per fruit. The second common green-white morph showed intermediate abundance and also showed leaky self-compatibility. Path analysis indicated that morph abundance had both direct and indirect effects on reproductive fitness, mediated by nectar, pollinator visitation and compatibility.

**Conclusions:** Reproductive success of the polymorphic *C. caulina* is a result of multi-trait interactions including pollinator interaction. That is, nectar traits and mating-system differences shape reproductive fitness of the morphs. We also highlight that in polymorphic perennial plants contemporary selection regimes may be acting alongside genotypic vestiges (historical genotypes present in extant population), thus complicating any frequency-dependent selection regime.

## Introduction

Floral color polymorphism (henceforth FCP) is the co-occurrence of two or more discrete color forms within a single population. The origins of FCP are usually explained by standard genetic processes such as mutation and recombination (Kellenberger et al., 2019; Stebbins, 2019), and the maintenance of morphs in any population is often explained by ecological selection such as by pollinators. Both vertebrate and non-vertebrate pollinators are well known for their use of floral color as a cue to locate and choose flowers resulting in pollinator-mediated selection shaping the success and decline of morphs within a population (Schiestl and Johnson, 2013; Rausher, 2008; de Jager and Ellis, 2014; Milano et al., 2016; Bigio et al., 2017; Dyer et al., 2021). However, there is mounting evidence that shows that floral color morphs also vary in other traits such as morphological, physiological, reproductive, and ecological traits, which can ultimately affect the morph’s fitness. Thus, recognizing the multivariate nature of FCP and characterizing how traits may covary among morphs is an essential precondition in order to identify selective agents that may be operating within a population.

Maintenance of FCP despite weak or absent pollinator preferences has been empirically shown (Schemske and Bierzychudek, 2007; Tang and Huang, 2010; Keasar et al., 2016; Hassa et al., 2020), and in several studies the target of selection within a population polymorphic for its floral and reproductive traits is not their floral color (Warren and Mackenzie, 2001; Subramaniam and Rausher, 2007; Dick et al., 2011; Fry and Rausher, 2017; Vaidya et al., 2018; Dafni et al., 2020; Table 1). Instead, these studies show that traits such as nectar rewards, flower size and number, pollen and ovule attributes, physiological tolerances, or mating-system differences alter reproductive outcomes independently of pollinator color choice and may play a greater role in the maintenance of the floral color morphs (Gomez, 2000; Wolfe, 1993; Carlson and Holsinger, 2013; Mu et al., 2011; Lau and Galloway, 2004; Kosi and Galloway, 2020; Arista et al., 2013; Ortiz et al., 2015). For example, seed viability differences linked to color morphs have been reported in *Boechera stricta* (Vaidya et al, 2018), while abiotic tolerance associated with color has been implicated in species such as *Parrya nudicaulis* and *Anemone coronaria* (Dick et al., 2011; Dafni et al., 2020). Some of the representative cases across plant families and selection regimes are summarized in table 1, which shows the growing body of examples that associate diverse functional traits with floral color morphs including life history strategies (annuals vs. perennials).

**Table 1:**
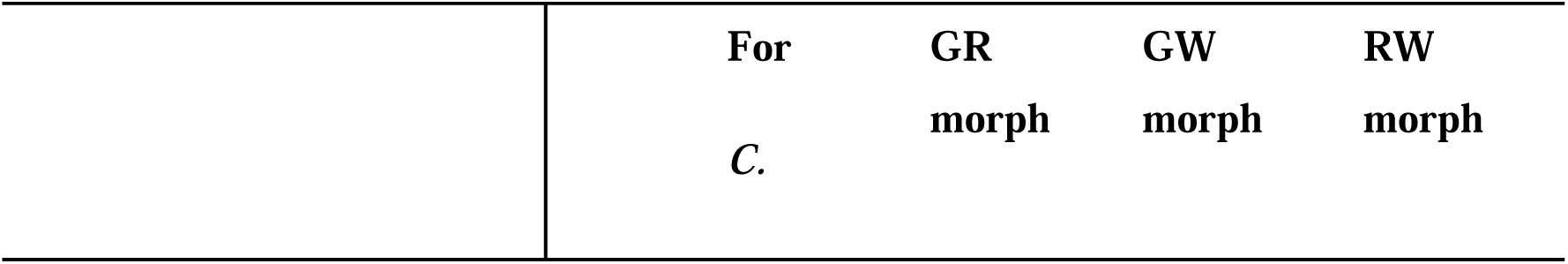
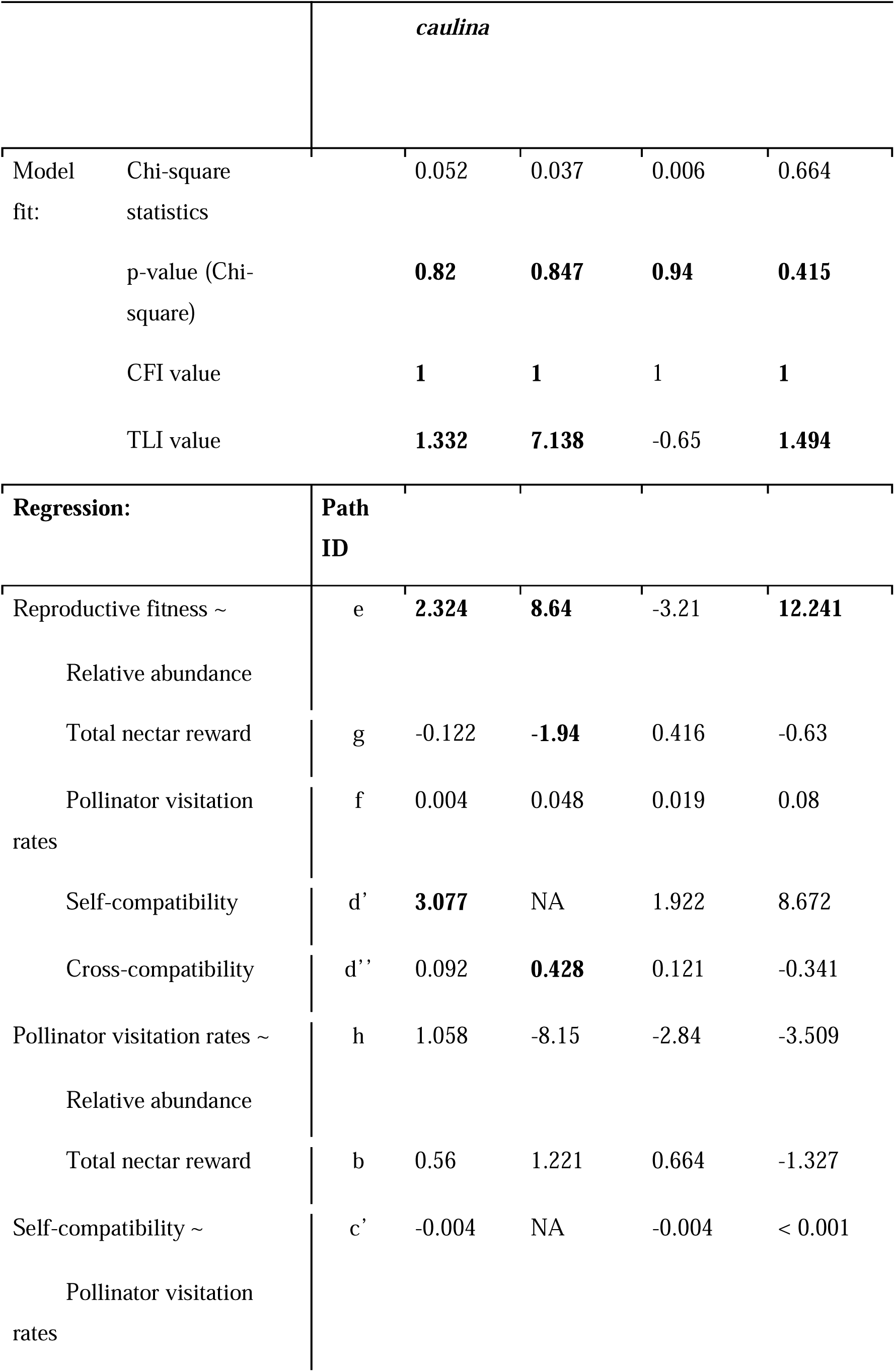
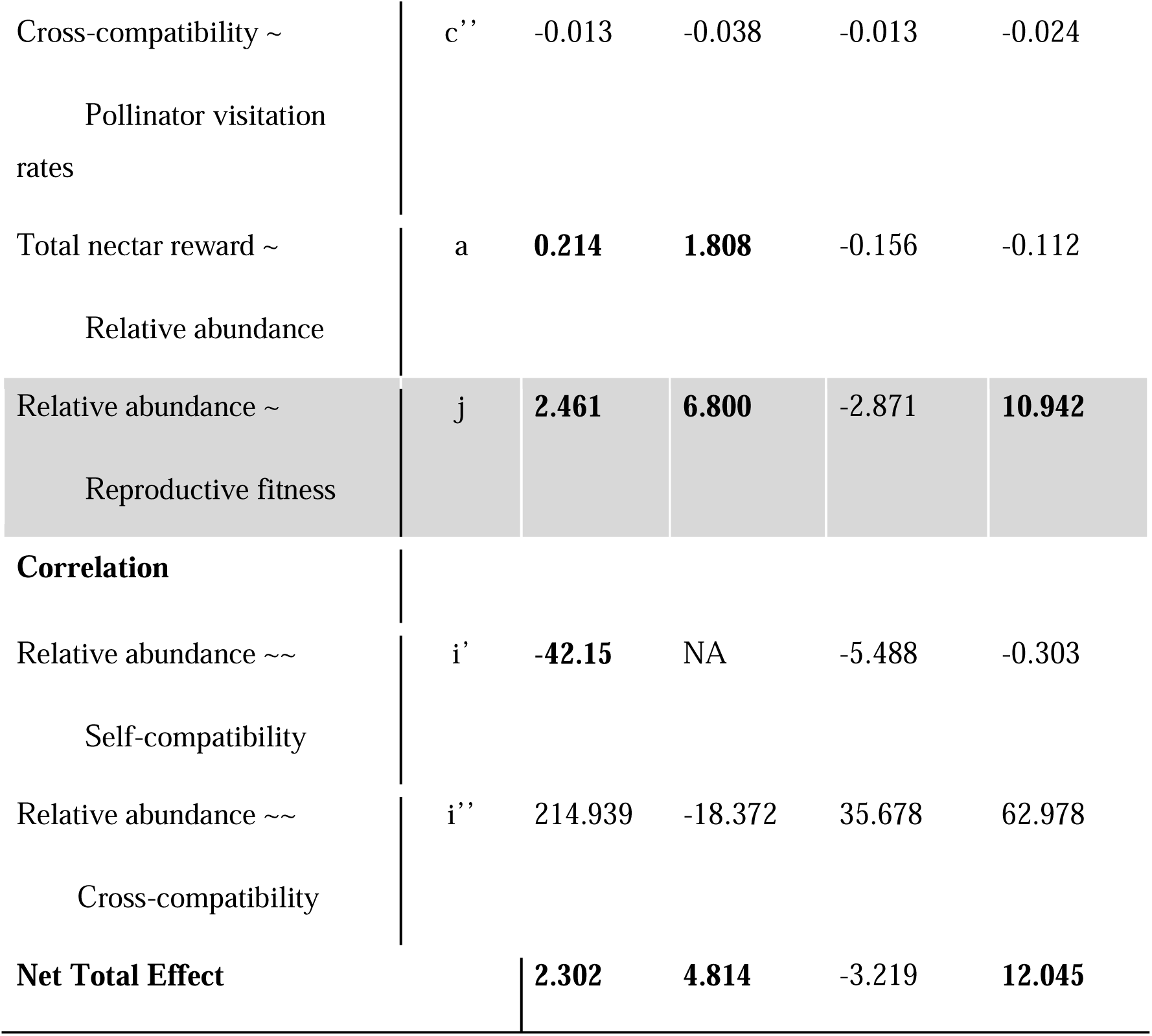
Results of path analysis for all four models. Bold values represent the statistical significance. Path IDs are the same as fig. 4.

The consequences of color polymorphism may differ across life-history strategies (Narbona et al., 2018; Sapir et al., 2021): in annuals, selection acts within a single reproductive cycle (Vaidya et al., 2018), whereas in long-lived perennials (rhizomatous and non-rhizomatous plants like trees) polymorphisms may persist or spread across multiple reproductive seasons (Snell and Aarssen, 2005; Friedman, 2020), allowing potential ecological or physiological advantages to accumulate over time. Therefore, life history strategy of a polymorphic species can further influence its morphological and genetic characteristics. Whereas frequency-dependent selection regimes may not even be detectable in perennials as past selections may persist in the current population as genotypic vestiges (historical genotypes present in extant population). This may be because change in morph frequencies often go undetected in perennial rhizomatous species that also propagate vegetatively (as reported in heterostylous species: Eckert and Barrett, 1993; Eckert et al., 1999; Barrett, 2015 and in dioecious species: Irish and Nelson, 1989; Petzold et al., 2013). Thus, clonality may buffer against rapid frequency shifts resulting in demographic feedback (Barrett, 2015). To the best of our knowledge there have been no studies that have looked into whether life history strategies influence the selection of polymorphism in perennial species. This is particularly striking for FCP, which is frequently reported in perennials but has not yet been empirically investigated.

Thus, plants with floral color polymorphism simplistically reveal at least the following three important points. First, color is often embedded in a broader phenotypic context; pigments such as anthocyanins have physiological functions (protection against UV, heat, drought, and pathogens) in addition to visual signaling (Narbona et al., 2018). Second, trait associations can alter selection indirectly: a morph that is less preferred by pollinators may nonetheless gain fitness if it enjoys higher autogamy, greater seed viability, or higher stress tolerance (Turelli et al., 2001; Schemske and Bierzychudek, 2007; Barazani et al., 2019; Vaidya et al., 2018). Third, perennial and clonal life histories complicate straightforward pollinator-mediated selection narratives. Clonal plants with long adult lifespans can accumulate historical demographic events which can decouple the effect of current selective agents on the extant morph frequencies.

Given these complex sets of drivers shaping a species that shows floral color polymorphism, we propose that as a first step it is critical to characterize phenotypic variations in perennial species that show FCP in order to understand extant selection regimes that may be maintaining the polymorphism within a population. To establish the variable traits among morphs, require documenting not just visual features such as color and pollinator visitation but also reproductive compatibility (autogamy, self-compatibility, cross-compatibility), floral rewards, other floral and vegetative morphological traits, and performance under abiotic stress. A multi-dimensional phenotypic characterization provides the empirical scaffolding needed to (a) identify which traits differ consistently among morphs, (b) evaluate the demographic and reproductive consequences of those differences, and (c) design targeted analysis of causal mechanisms.

Here, we explore a polymorphic, nocturnal ginger *Curcuma caulina* from the family Zingiberaceae. All gingers are perennials due to their rhizomatic habit and they offer ideal grounds to test the role and maintenance of polymorphism in floral forms. Several genera show pronounced intraspecific variation in flower and bract colors, and within the Asian tropics a range of polymorphisms (stylar, bract color, and floral color polymorphisms) has been reported across genera such as *Alpinia*, *Amomum*, *Curcuma*, *Hedychium* and *Roscoea* (Poulsen & Lock, 1997; Harris et al., 2000; Leong-Skornicková et al., 2007; Mood et al., 2013; Yadav, 2024; Yadav and Gowda, 2024). In some taxa, floral polymorphism has been linked to pollinator shifts and speciation dynamics (Takano et al., 2009; Paudel et al., 2019), but detailed, multi-trait descriptions remain rare for many species.

In this study we present a comprehensive characterization of bract-color polymorphism in *C. caulina*, which is endemic to the laterite-plateaus of the Northern Western Ghats (Figure S1). In our preliminary observations we had noted that morphs showed variable abundances within a population, with a few dominant, and several rare morphs. Due to the absence of any prior ecological studies on *C. caulina*, we focused on three complementary trait axes in order to characterize intraspecific morph variations. They are: morphology (bract and floral display), pollinator visitation, and reproductive compatibility traits (autogamy, cross-compatibility, and seed set). These traits form the immediate causal chain linking phenotype to reproductive output in animal-pollinated plants: bract color and rewards influence pollinator attraction and visitation; visitation determines pollen transfer; and reproductive compatibility converts pollen receipt into realized fitness. These axes capture both pre-pollination (pollinator attraction and pollen receipt) and post-pollination (fertilization and compatibility) processes and can be quantified in the field. More generally, the questions we ask are ecological. These traits are analogous to those examined in studies investigating interspecific competition; however, we utilize them within a polymorphic species that exhibits intraspecific variations among its morphs.

Using field experiments we predicted that since higher nectar reward is known to increase visits by pollinators, morphs with better nectar rewards will show higher pollinator visitation rates, higher pollen transfer, and increased fruit set and seed set. Since we had noted differences in morph abundances within our study site, we predicted that the most common morph will show the highest nectar rewards, visitation rates, and fruit set and seed sets. We also expected that rare morphs may compete for pollinator visits with the other more abundant morphs, resulting in a trade-off between morph abundance and reproductive compatibility.

Based on these expectations, we addressed the following questions:

i. Do nectar rewards, reproductive compatibility, and pollinator visitations differ among the color morphs?
ii. Do pollinator and plant reproductive traits explain reproductive success and do these correlate with the observed morph abundance?

## Materials and Methods

### Study species and study site

*Curcuma caulina* (Zingiberaceae) is a rhizomatous, mass-flowering, perennial herb that flowers between July to October (Fig. S1). The plant is 30–50 cm tall with a single shoot bearing a terminal inflorescence which consists of 10-18 fertile ‘lateral bracts’ that bear flowers (Fig. S1). Flowers are born in cincinni (1–3 flowers per cincinnus), are nocturnal (anthesis at 18:00), and last 12–14 hours only. The stigma is funnel-shaped and projects beyond the anther, and fruits mature within the bracts. *C. caulina* is endemic to the northern Western Ghats of India and this study was carried out on Kaas Plateau (कास पठार; 17°43’12.36” N, 73°49’28.0956” E, elevation 1225m, Fig. S1), Satara, Maharashtra, India between 2019 and 2022. Kaas Plateau is an elevated ferricrete outcrop (red laterite crust) and a UNESCO Natural World Heritage site recognized for its floral diversity and high endemicity (Shrotri et al., 2025).

### Bract color polymorphism and variation in floral morphology and nectar rewards

The color of the lateral bracts of *C. caulina* individuals varied within the population from predominantly green to predominantly red colored inflorescence (Fig. S2a). To determine the frequency of bract color morphs, we identified and counted all morphs within randomly placed ten transects, each measuring 10 x 4 m^2^. Variations in the color of the lateral bracts were first identified based on the extent of the reddish-pink coloration (0% to 100%) on the first or basal lateral bract. This color difference was visually identified and also quantified using handheld spectrophotometer (Ocean Optics; Fig. S2a, b).

To morphologically characterize the morphs, we measured a total of 22 vegetative and floral characters using digital vernier caliper and rulers (n=30 per morph; character list in Table S1). To assess morphological variations among the morphs, we ran a non-metric multidimensional scaling (nMDS) analysis with Bray–Curtis dissimilarity using the VEGAN package (Dixon, 2003) in R (4.3.0; R core Team, 2021). We used the first three dimensions, to construct three-dimensional (3D) plots, which were used to visualize both intra-morph and inter-morph variations. The statistical difference among the overlapping clusters (morphs) was measured using the ANOSIM test, where the Bray–Curtis dissimilarity matrix with 1000 permutations was used (VEGAN package; Dixon, 2003).

We measured nectar volume and concentration in 106 bagged flowers (32 to 38 unopened buds per morph) between 18:00 and 18:30 hrs by destructive sampling. The nectar volume (μl), concentration (% sucrose), and nectar energy (cal) were compared among the three morphs. Nectar volume was measured using calibrated glass microcapillary tubes (Drummond Scientific Company), and the sucrose concentration in nectar was determined using a handheld refractometer (ERMA BRIX 0∼55%). The brix sucrose % value was then converted into sugar (mg) per 1 ml of nectar using the table given in the CRC Handbook of Chemistry and Physics (Lide, 2004) and as outlined by Bolten *et al*. (1979). This value was used to calculate the amount of sugar in a given volume of nectar, and it was converted into calories (1 mg of sucrose = 4 calories as given in Heinrich, 1975). Differences between the morphs were tested using the Kruskal-Wallis test followed by Dunn’s test as a post-hoc; the significance was set at *p*<0.05; analysis was carried out in R (4.3.0; R Core Team, 2023).

### Natural pollinator visitations

To determine if pollinators exhibit differential visitation towards a particular morph, pollinator visits were monitored by night vision CCTV cameras (Ae Zone CCTV IR day/night vision USB camera) between 18:00 hrs and 06:00 hrs. These observations were carried out in 18 randomly selected observation plots on the plateau, where each plot consisted of three to six individuals. All morphs were observed for a total of 648 hrs (i.e., 216 hrs per morph). Cameras were positioned approximately 1 to 1.5 m away from the focal plants such that all inflorescences were visible in their entirety in the field of view. The recordings were manually scored, where the total number of visits by each pollinator and the total number of flowers present in the plot were measured. We calculated the pollinator visitation rate as the total number of visits per flower per hour for each morph. The visitation rates were summed for every hour between 18:00 to 06:00 hrs in order to record the temporal shift in pollinator visitation rate.

Natural pollinator visitation rates (number of visits per flower per hour) were compared among the morphs on a temporal scale using the circular statistical analysis in ORIANA (v. 4.02 Kovach Computing Services; Kovach, 2011). Temporal differences in visitation rates between the morphs were tested using the Mardia-Watson Wheeler test (or uniform score test; Zar, 1999). Since the pollinator observation plots were not always located within the morph-census transects, the pollinator visitation rates were standardized for morph frequency using Monte Carlo Simulation (n = 10,000). The morph frequency and pollinator visitation rates for each morph were utilized to estimate the distribution of their product for the respective morphs. To test for inter-morph differences in pollinator visitation rates we used the peak visitation rates from 19:00 hrs to 22:00 hrs, which accounted for ∼85% of all pollinator visits. We used the Kruskal-Wallis test followed by Dunn’s test as a post-hoc test at *p*<0.05. All statistical analyses were carried out in R (4.3.0; R Core Team, 2023).

### Physiological compatibility and natural fruit set

To comprehensively evaluate the physiological self-compatibility and cross-compatibility rates in the three floral color morphs, we carried out a total of 15 hand-pollination treatments. Since we identified three bract color morphs in this population (refer to results for within population morph variability), reproductive compatibility was tested for these three morphs, where one morph was rare and two were common. Each morph was treated separately, and the pollination treatments included three autogamy, three self-pollination, and nine cross-pollinations that included three intra-morph cross-pollinations and six inter-morph cross-pollinations were the identity of a morph as the male or female parent was recorded.

Autogamy treatment included unmanipulated, bagged flowers that were checked for fruit set after four weeks. For cross-pollination treatments, flowers were emasculated and mixed pollen grains from 10-15 individuals were manually transferred on marked flowers. Flowers were observed for fruit set in ∼4 weeks and percent success rate in reproduction was measured for each treatment using the number of fruits per treatment as well as the number of seeds per fruit (seed count). We statistically tested the reproductive success rates between self- and cross-pollination treatments for each morph and compared male versus female reproductive success for the morphs using Fisher’s exact tests. Comparative analyses were also carried out for the seed counts from self- and cross-pollination treatments using the Kruskal-Wallis test followed by Dunn’s post-hoc test with significance set at *p* < 0.05.

We quantified the reproductive success of natural pollination events within the study population at the end of the flowering season in mid-October by measuring fruit set in ≥ 500 individuals per morph and seed counts in ≥ 50 per morph. For each inflorescence, we estimated the total number of flowers per inflorescence by multiplying the number of lateral bracts with the number of flowers estimated in a cincinnus. Fruit-set per inflorescence was then calculated as the ratio of the number of matured fruits to the total number of flowers estimated for a particular inflorescence. We used Kruskal-Wallis test to compare both natural fruit set and natural seed count among morphs. Dunn’s post-hoc test was used with a significance set at *p* < 0.05.

### Path analysis to test traits effects on the reproductive fitness of the morphs

We used path analysis to understand the direct and indirect effects of abundance of a morph on its natural fitness as described by Mitchell (1993). Path analysis was tested for the entire Kaas population of *C. caulina* as well as for each morph independently (n = 300 individuals; i.e., 100 per morph) in order to assess the net effect and important selection regimes within the population and within each morph. Three indirect effects included in the path analyses were-total nectar reward, pollinator visitation rates, and physiological compatibilities. Values for the total nectar reward were normalized with morph abundance and of natural fitness with seed-set using Monte Carlo simulations (n = 10,000). Path analysis was performed in R (4.3.0;

R Core Team, 2023) using the *lavaan* package (Rosseel, 2012) and path estimates were computed using 10,000 iterations in the bootstrapping procedure. Estimates of model fit were computed using Chi-squared goodness of fit, Comparative Fit Index (CFI) and Tucker-Lewis Index (TLI). A nonsignificant chi-squared value indicates that a model is not significantly different from the observed correlations in the data and is therefore a good fit whereas CFI and TLI indices above 0.9 are considered to be a good fit (Mitchell, 2001). Finally, we tested the hypothetical relationship between reproductive fitness and relative abundance using a regression model which was not part of the main path analysis.

## Results

### Floral (bract) color polymorphism and morphological variability

We identified six bract color variants with an almost continuous distribution within the population (Fig. S2c). However, using Ford’s criteria for polymorphic forms (Ford, 1945; Fig. S2), we chose three most common variants (0% red, 60% red, and 100% red; henceforth referred to as ‘morphs’) in this study. These three morphs were named based on the coloration of their lateral and coma bracts as follows: the 0% variant is Green-White morph (GW), the 60% variant is Green-Red morph (GR), and the 100% variant is Red-White morph (RW; Supp Fig. 1a). The distribution of these morphs was unequal, with GR morphs being the most common (∼ 44% occurrence), followed by GW (∼ 29%) and RW (13%). The remaining three bract color variants (15%, 30%, and 45%) were extremely rare, each constituting less than 5% of the population (Fig. S2c).

**Figure 1.**
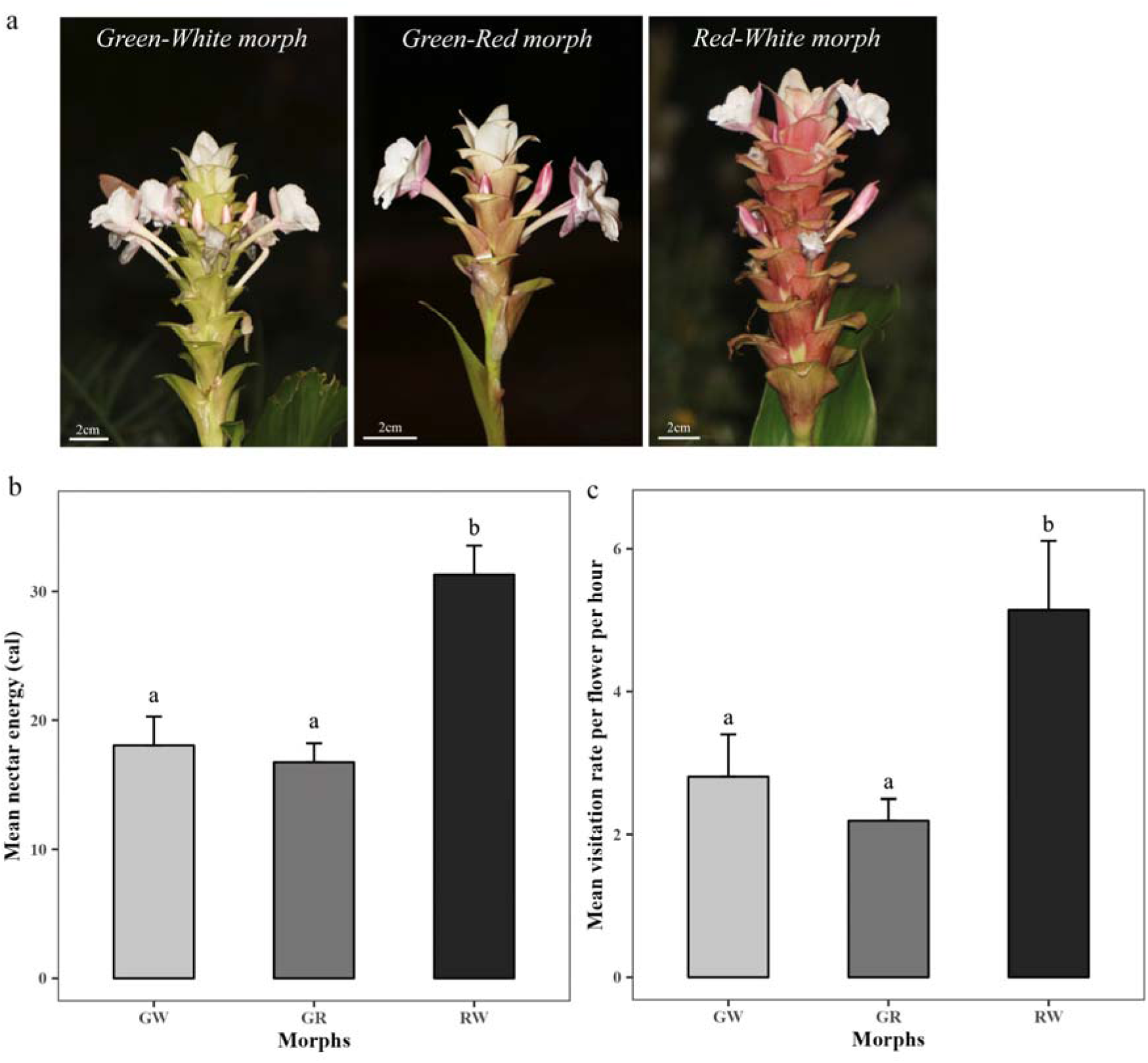
Three bract color morphs of *C. caulina*, their nectar rewards energy, and natural pollinator visitation rates. a. From left, GW, GR, and RW morphs of *C. caulina*, b. The mean (± s.e.) nectar energy (cal), c. Bar plot represents pollinator visitation rates (mean ± s.e.) measured for each morph in the observation plots (y-axis) and simulated pollinator visitation rates (mean ± s.e.) calculated for the population shown by red dots. Different alphabets indicate significant differences between morphs (*p* < 0.05).

The abundance of the three morphs differed within the population. The GR morph (98.6 ± 7.2 individuals; ∼44 %) and GW morph (75.7 ± 7.2 individuals; ∼29 %) were the most common morphs; their frequencies were not significantly different from each other (Kruskal-Wallis X2 = - 0.692, df = 1, *p* = 0.2446). The rarest of the three morphs was the Red-White morph (RW) with 24.2 ± 2.0 individuals (∼13 %) in the population, and the RW morph had significantly fewer individuals when compared to both GR and GW (RW vs GR, Kruskal-Wallis X2 = - 2.427, df = 1, *p* = 0.0076 and RW vs GW, Kruskal-Wallis X2 = 1.736, df = 1, *p* = 0.0413 respectively). All other bract color variants totaled to ∼14% in the population and thus were extremely rare. The non-metric multidimensional scaling (nMDS) analysis of the 22 floral and vegetative morphometric characters (Table S1; Fig. S2) failed to form distinct clusters separating the morphs by their morphological traits. Further, the Analysis of Similarities (ANOSIM) test (*p* = 0.4835) and the associated ANOSIM R-value (ANOSIM R = - 0.0038) also indicated no statistical significance in the differentiation of these clusters for each morph. Since we detected polymorphism in bract colors and not floral colors, henceforth, the term color polymorphism refers to only bract color polymorphism in *C. caulina*.

### Variability in nectar traits and hawkmoth visitation rates

Nectar is the primary pollinator reward in *C. caulina*, and there was no significant difference between the total nectar rewards (assessed in calories as mean ± s.e.) of the two common morphs GR (16.74 ± 1.47) and GW (18.04 ± 2.2). However, the total nectar reward was highest for the rare RW morph (31.31 ± 2.25) due to both its higher nectar volume and concentration (Fig. 1b; Table S2; Kruskal-Wallis X2 = 33.53, df = 2, *p* < 0.0001). Flowers of *C. caulina* were visited by two species of hawkmoths with long probosces: *Agrius convolvuli* (∼12.4 cm) and *Hippotion rafflesii* (∼4.6 cm, Fig. S3; Sphingidae). During foraging, the moths insert their proboscis into the floral tube, hover over the flower, and the sticky pollen is carried on the proboscis of *A. convolvuli* (Fig. S3) or on the head in the case of *H. rafflesii*. We did not observe the curling of proboscises during foraging, as noted by Smith et al. (2022). The average time spent by each pollinator per flower was ∼ 1.5 seconds for *A. convolvuli* and ∼ 2.5 seconds for *H. rafflesii*, and the rupture in anther theca could be used as a visual feature of flowers visited by hawkmoths.

The circular statistical analysis of visitation rates across the anthesis time showed no temporal differences in pollinator visitations among the three morphs (Rayleigh Test; *p* < 0.0001; Fig. S4). The mean pollinator visitation time, denoted by the circular mean vector, µ was noted to be between 21:00 to 22:00 for all morphs (Fig. S4). All three morphs showed the highest hawkmoth activity during the first few hours of anthesis, between 19:00 hrs to 23:00 hrs, which gradually decreased until there was no activity by 06:00 hrs the following morning (Fig. S4). Pollinator visitation rates of the three morphs during the peak visitation hours were statistically different from each other with RW morph showing the highest visitation rate (mean number of visits per flower per hour ± s.e.; 5.14 ± 0.96), followed by the GW morph (2.8 ± 0.59), and GR morph (2.19 ± 0.3; Fig. 2c; Kruskal-Wallis chi-square = 6.52; df = 2; *p* = 0.03839; Table S3). However, the visitation rate standardized for morph abundance showed a significantly lower visitation rate (Kruskal-Wallis chi-square = 3.77; df = 2; *p* = 0.01511) to the RW morph (mean number of visits per hour ± s.e.; 119.94 ± 10.75), than the common GR (175.15 ±17.29) and GW morphs (171.39 ± 18.72; Fig. 1d;).

**Figure 2.**
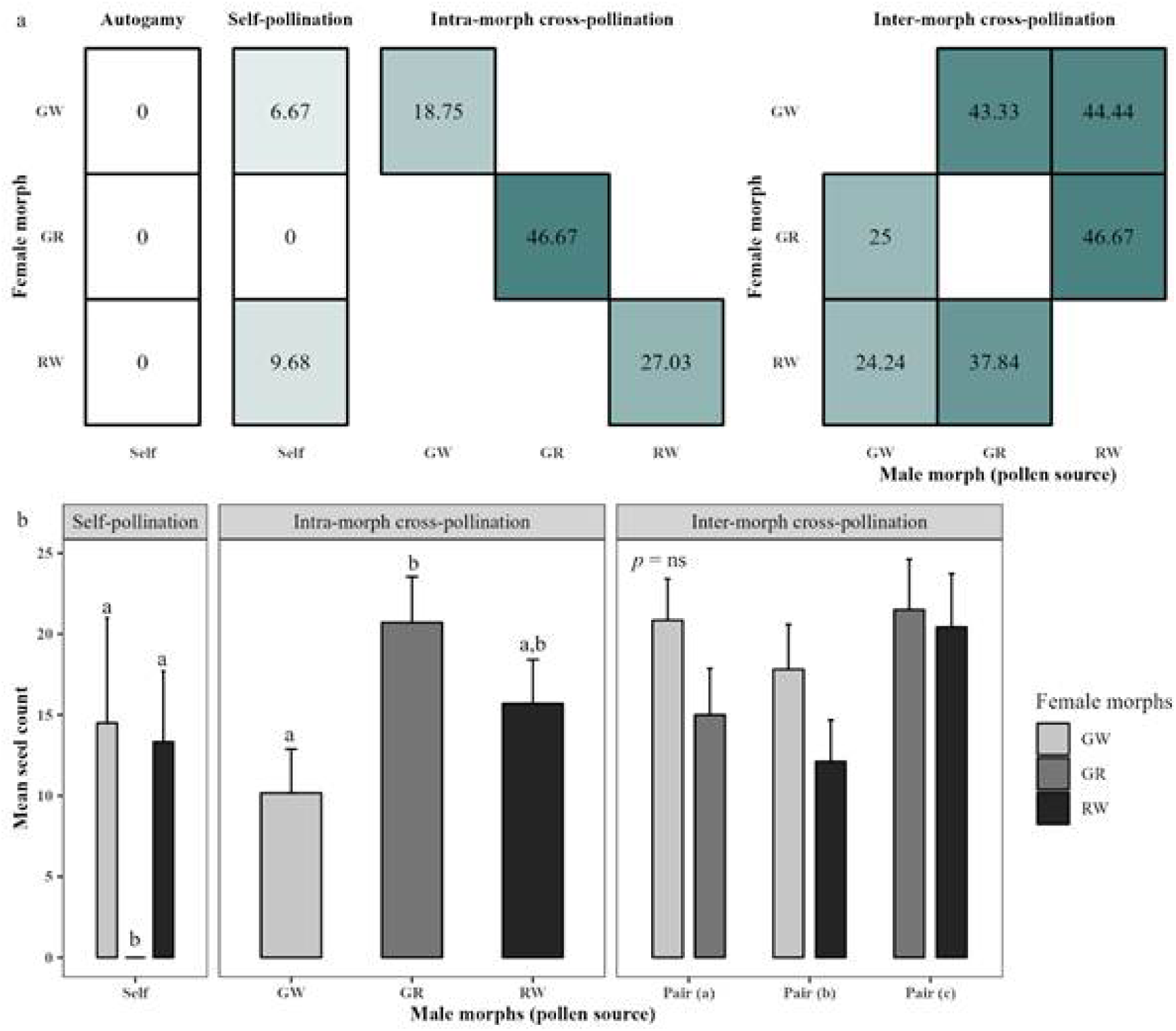
Physiological compatibility between the three floral color morphs of *C. caulina*. a. Heatmap illustrates the percent success rate (values inside box) from 478 hand-pollination from 15 treatments. The male morphs (pollen source) are displayed on the x-axis, while the female morphs (stigma identity) are shown on the y-axis. b. Mean (± s.e.) seed count from the pollination treatments in the three morphs of *C. caulina*. The male donors (pollen source) are displayed on the x-axis while the boxes represent female recipients (stigma identity). Significant differences are shown by different alphabets.

From video recordings of pollinator visitations, we also observed that hawkmoths revisited the same flowers within an RW morph (∼ 10-12 times), once they discovered this rare but nectar-rich inflorescence. In contrast, the two common morphs GR and GW with low nectar rewards received lower visitation rates, and pollinators rarely revisited the flowers. Thus within-inflorescence visits were higher in the rare RW morph compared to the common GR and GW morphs.

### Physiological compatibility and natural fitness vary among the morphs

A total of 471 hand pollinations were carried out (with n ≥ 30 per treatment) where the three intra-morph cross-pollination treatments were: GW x GW, GR x GR, and RW x RW (male x female morph) and will be interchangeably referred to as ‘cross-pollination’ treatments. Inter-morph cross-pollination treatments were categorized into three pairs to investigate the effect of male and female parents in each treatment. These crosses are (male x female) identified as pair a: GW x GR and GR x GW, pair b: GW x RW and RW x GW, and pair c: GR x RW and RW x GR. Out of 471 hand-pollination treatments, only 108 treatments set fruits (Fig. 2a). None of the autogamous treatments set fruits. Selfing treatments failed to set fruits exclusively in the GR morph, whereas the GW and RW morphs demonstrated leaky self-compatibility with success rates of 6.67% and 9.68%, respectively. Fruit-set success in selfed treatments was 5.8%, which was significantly lower (χ2 = 31.299, *p* < 0.0001; Table S4) when compared with fruit-set success in the intra-morph cross (48.38%) and inter-morph cross (58.4%). The GR morph demonstrated the highest fruit set in the intra-morph outcrossed treatment with a success rate of 46.67%, followed by the RW morph at 27.03% and the GW morph at 18.75% (Fig. 2a).

In the inter-morph crosses, we noted donor-specific differences in the GW and RW morphs. The donor specificity was related to whether a morph was used as a male parent or a female parent in the inter-morph cross-pollination treatments. In crosses where GW morph was the male parent (i.e., pollen donor), success rates were significantly lower (pair a: GW x GR, 24.24% and in pair b: GW x RW, 25%) when compared to crosses where it was used as a female parent, that is pollen recipient (in pair a: GR x GW, 43.33% and in pair b: RW x GW, 44.44%, Fig. 2a, Fisher’s exact test *p* = 0.027, odd ratio = 0.419). Whereas when RW acted as the male parent (pollen donor) in inter-morph crosses, fruit set success was higher (pair b: RW x GW = 44.44%; pair c: RW x GR = 46.67%) compared to when it served as the female parent (pair b: GW x RW = 25%; pair c: GR x RW = 37.84%) however, Fisher’s exact test did not yield the significant difference (*p* = 0.1129, Odd ratio = 1.8).

Among all hand-pollination treatments, self-pollination treatments exhibited the lowest seed count (mean number of seeds per matured fruit ± s.e.; 8.63 ± 3.15), followed by intra-morph crosses (16.93 ± 1.8) and then inter-morph crosses (18.63 ± 1.2). In the self-pollination treatment, the GR morph exhibited a significantly lower seed count (zero) compared to the GW morph (14.5 ± 6.5) and RW morph (13.33 ± 4.37; Kruskal-Willis X2 = 5.2499, df = 2, *p* = 0.07244; Fig. 2b; Table S5 and S6). In the intra-morph cross-pollination treatment, the seed counts varied significantly among the morphs. The GW morph exhibited the lowest seed count (10.72 ± 2.7) compared to the GR morph (20.71 ± 2.8) and the RW morph (15.7 ± 2.7; Kruskal-Willis X2 = 5.2499, df = 2, *p* = 0.07244; Fig. 2b; Table S6). Seed count in inter-morph cross-pollination treatments was not significantly different among the three pairs (Fig. 2b; Table S6). Finally, the mean total fruit set per plant in open pollination treatments was recorded to be equal in all three morphs (Fig. 3a). However, the mean seed count per fruit was significantly different among the morphs (Kruskal-Wallis X2 = 36.3222, df = 2, *p* < 0.0001) with the GR morph being the highest (23.7 ± 1.5), followed by GW (20.3 ± 1.2) and RW (11.5 ± 1.4; Fig. 3b). We also noted that the seed counts recorded in the open (natural) pollination and self-pollination treatments of RW were similar (Fig. 2a and 3a).

**Figure 3.**
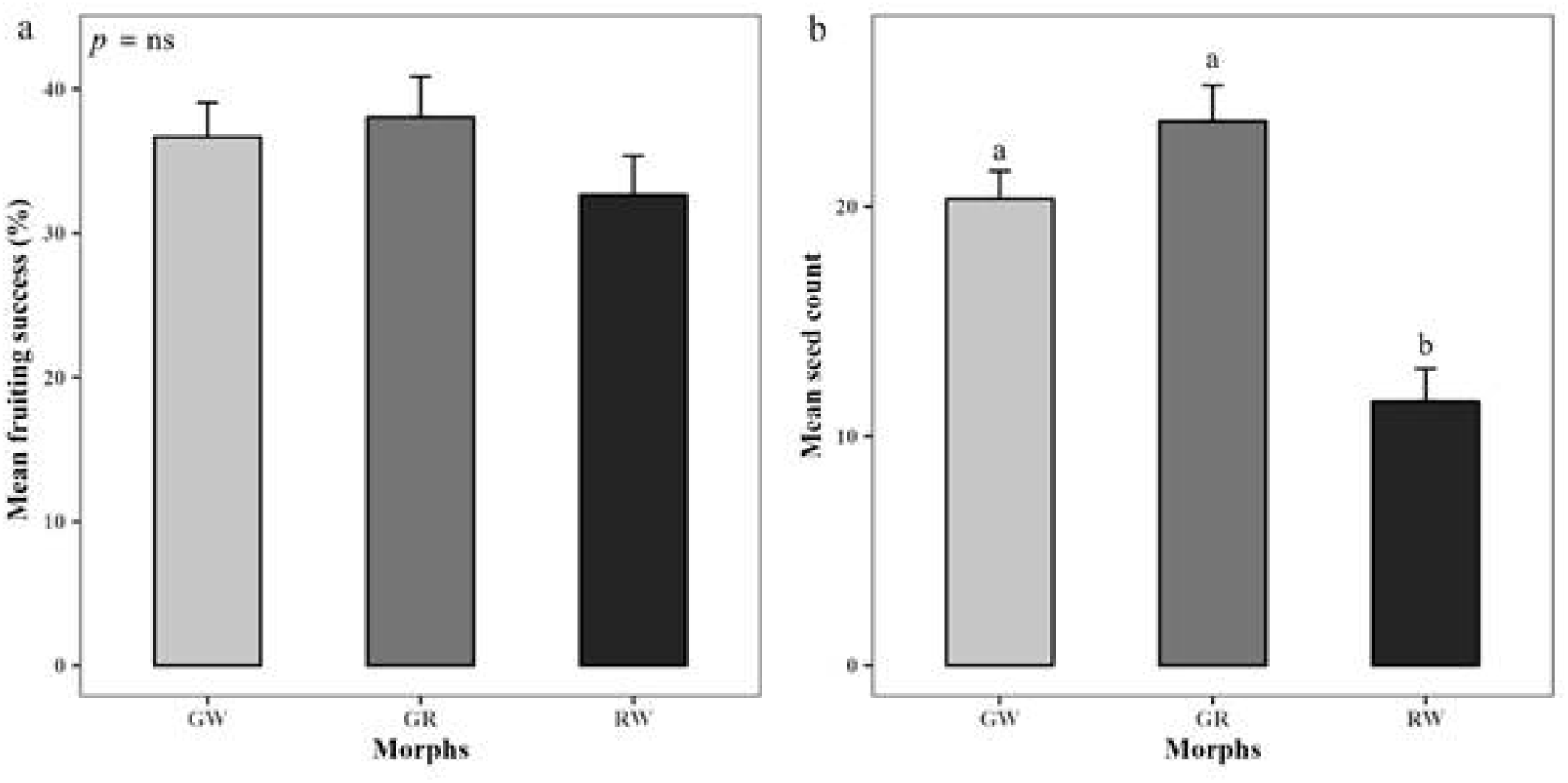
Natural fitness of the three morphs of *C. caulina* measured as A. mean (± s.e.) reproductive success (% success) of open pollination treatments and B. mean (± s.e.) seed count of the successful fruits from those open pollination treatments. Small letters indicate significant differences between female morphs. Results of the statistical test between morphs are given in Table S6.

### Associating morph abundance and floral traits

In the path analyses, the five variables resulted in one direct path between abundance and reproductive fitness (e), seven indirect paths (a to d and f to h), one correlation path between abundance and type of compatibility (i), and one path on the effect of reproductive fitness on relative abundance (j; Fig. 4). The Chi-squared goodness of fit test, CFI, and TLI scores identified a good fit for all four models (*C. caulina*, GR, GW and RW morph; Table 1). In the case of the GW morph, the model fit as estimated by TLI values was not good (TLI = -0.65), and we did not find any significant relationships between the traits as well as for the net effect. We found that both the direct effect (path e in Fig. 4) and the net effect of morph abundance on reproductive fitness were positive and statistically significant in three (*C. caulina*, GR morph, and RW morph) of the four path analyses models (Table 1).

**Figure 4.**
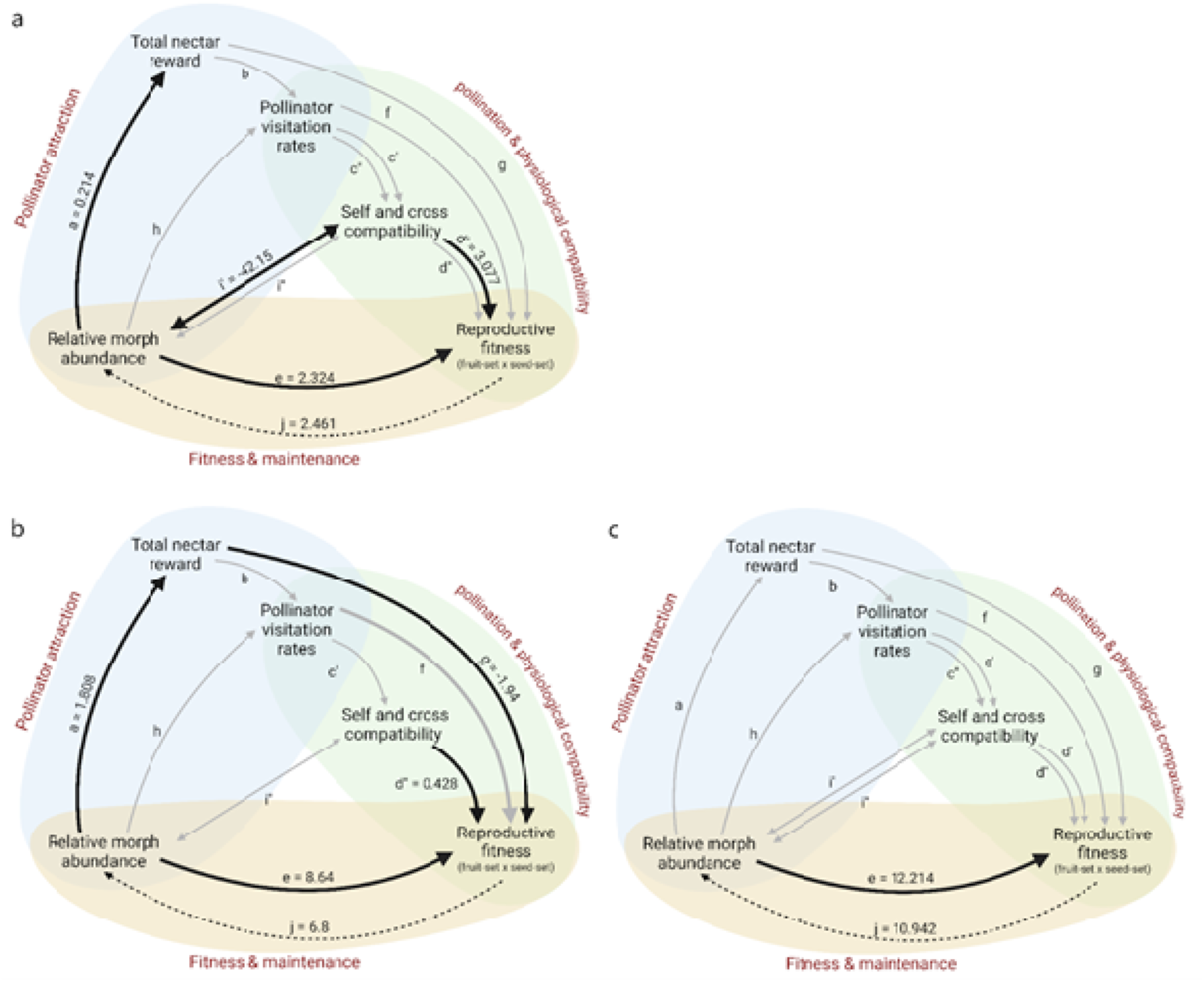
Path diagrams illustrating the effect of morph abundance on reproductive fitness, analysed a) as species-specific model, b) for the common GR morph and c) for the rare RW morph. One-headed arrows (paths a—h and j) indicate regression, while the two-headed arrow (path i) indicates correlation. Thick arrows denote significant effects (P < 0.05), with the strength of regression or correlation specified adjacent to them. Path IDs for all direct and indirect effects are provided, and their strengths can be referenced in Table 1.

In the species-specific model, *C. caulina* in KAS showed a positive and significant effect of only morph abundance on total nectar reward (a), self-compatibility on reproductive fitness (d’), and finally, reproductive fitness on relative morph abundance (j). The correlation between morph abundance and self-compatibility was also noted to be positive (i’’). For the within-morph path analyses, significant effects among indirect effects were observed only in the GR morph, although positive effects were present across several paths for the other two morph-specific models (GW and RW). In the GR morph, the following paths were significant - morph abundance and total nectar reward (a), total nectar reward and reproductive fitness (g), cross-compatibility and reproductive fitness (d’’), and the regression of reproductive fitness on morph abundance (j). The direct effect between relative morph abundance and reproductive fitness (e), the regression of the reverse effect, that is, between reproductive fitness and abundance (j) was significant and positive in RW but negative and not significant for GW. We failed to detect any significant relationships between the other traits in the RW morph, although notably, the effect of self-compatibility on reproductive fitness was positive, whereas the effect of cross-compatibility on reproductive fitness was negative but not significant (path d; Table 1).

## Discussion

Floral color polymorphism, variable morph abundances, and population shifts in morph frequencies are well-documented across angiosperms. However, only few studies have explored the breadth of phenotypic landscape of FCP, that is, how floral color covaries with morphology, physiology, mating systems and ecological interactions, and this is particularly rare for long-lived, clonal taxa (Elam & Linhart, 1988; Lau & Galloway, 2004; Caruso *et al*., 2010; Berardi *et al*., 2016; Koski *et al*., 2020; Jacquemyn & Brys, 2020). Our study of bract-color morphs in the nocturnal ginger *Curcuma caulina* illustrates why such breadth matters. On lateritic plateaus which are known for a strongly seasonal environment, abiotic stressors are pronounced. We found that morph identity is tied not only to differential nocturnal visitation by hawkmoths but also to nectar investment and mating-system differences that together determine realised reproductive outcomes. Interpreting these patterns therefore requires an integrated view that accounts both for immediate pollinator behaviour and for longer-term demographic and physiological dynamics characteristic of perennial, clonal species.

Floral color variation is widespread, and in tropical lineages such as the order Zingiberales and within the family Zingiberaceae many species exhibit striking floral and bract color variation (Maas-Van De Kamer et al., 2016; Temeles et al., 2000; Gowda & Kress, 2013; Leong-Skornicková et al., 2007; Saryan et al., 2020). Previous studies have suggested that specialisation to particular pollinators may drive morph differentiation. At one extreme of this idea lies the ‘one pollinator–one morph’ scenario (Temeles et al., 2013; Meléndez-Ackerman et al., 2005) that is empirically reported in the Caribbean *Heliconia caribaea* (Temeles et al., 2013) and trans-Himalayan *Roscoea* spp. (Paudel et al., 2019). And the other extreme is, one pollinator-multi morphs and multi pollinator-multi morphs interactions reported in the Caribbean *Heliconia* spp. (Martén-Rodríguez et al. 2011; Gowda et al., 2012; Gowda and Kress, 2013). To understand the role of pollinators in shaping morph frequencies in a polymorphic species we addressed whether pollinator specialisation offers a satisfactory explanation in the case of *C. caulina*. Within the Kaas population all morphs displayed classic moth-pollination syndromes (Johnson *et al*., 2017), such as nocturnal opening, white colored flowers, and long floral tube with copious nectar. True to its syndrome, we discovered that *C. caulina* is exclusively pollinated by nocturnal hawkmoths and the moths show differential visitation towards the morphs. However, in our study, pollinator identity and visitation patterns alone did not provide a full account of morph-level fitness. Like many classic pollinator-mediated models we expected morphological differences (tube length, flower size, display) to drive pollinator sorting and hence differential fitness (Meléndez[:Ackerman and Campbell 1998; Gomez 2000; Majetic et al. 2007; Milano et al. 2016); interestingly, in *C. caulina* those morphological traits do not differ among morphs, and instead only nectar investment and mating-system differences were the two most defining axes. Thus we did not see the strict ‘one pollinator–one morph’ outcome in *C. caulina*. We propose that the trait space through which the hawkmoths discriminate and affect the plant’s fitness appears to rely primarily on floral reward rather than due to gross morphological matching.

In *C. caulina*, the first ecological trait that showed direct effect on reproductive fitness is morph abundance (path e, Fig. 4). The rocky lateritic plateaus of Northern Western Ghats are known for their extreme weather conditions suggesting the presence of strong abiotic selection factors (Watve, 2013; Shrotri et al., 2025). Our identification of the tri-modally distributed morphs with distinct physiological features suggests that these morphs may have been under different selection pressures in the past. For example, anthocyanin-rich red bracts of RW morph may confer UV or drought tolerance (Berardi et al., 2016) which may have been advantageous in the rocky laterite habitats like Kaas. Unequal morph abundance has been commonly reported in plants showing FCP. Narbona et al. (2018) noted that more than 80 % of Mediterranean species with FCP exhibit skewed morph distributions that often fluctuate spatio-temporally due to abiotic pressures (Dormont et al., 2010; Tang et al., 2016). Thus, we propose that abiotic adaptations, along with vegetative propagation, could decouple morph frequencies from contemporary selection, allowing relic traits to persist (Barrett, 2015; Záveská et al., 2012) resulting in a polymorphic population with a few rare but persistent morphs, as noted in *C. caulina* at Kaas plateau.

Physiological compatibility is the second trait that we identified as affecting the plants reproductive fitness (path d). While physiological traits are not expected to be different between morphs, we noted two different reproductive outcomes in two separate morphs of *C. caulina* - a) improved reproductive fitness via strong self-incompatibility in the most common GR morph, and b) self-compatibility providing reproductive assurance in the rare RW morph. The second strategy has been suggested theoretically (Lloyd 1992; Holsinger, 2000) and empirically in *Ipomea purpurea* (Subramaniam and Rusher, 2000; Fry and Rausher, 2017), and *Lysimachia arvensis* (Arista et al., 2013; Ortiz et al., 2015). However, our results demonstrate that contrary to our expectations reproductive strategies can vary significantly both within a species and within a population.

The third trait that showed a direct effect on reproductive fitness was floral nectar. The absence of both high nectar reward and higher pollinator visitations translating to higher reproductive fitness (fruit set and seed set) suggests that the rare morph may have a trade-off between its nectar investment and observed female reproductive success. This is in contrast to the pattern noted in *Ipomoea purpurea* where rare, nectar-rich morphs in a population benefit from higher pollen export (Fry & Rausher, 1997). We speculate that the higher nectar rewards in the rare RW morph may boost pollen export as noted in *Antirrhinum* sp. (Jones & Reithel 2001) and therefore may be maintained in the population despite the observed trade-off.

So, what factors explain the observed disparity in morph abundances in *C. caulina* at Kaas? We propose a multi-trait hypothesis involving an interplay between nectar reward, pollinator attraction, and physiological compatibility (self vs. cross), which collectively influence the reproductive success of different morphs and subsequently affects their population abundance. The common morphs show higher reproductive success despite having lower nectar energy probably because their self-incompatibility ensures that all seeds are outcrossed and with higher seed-set (fitness). On the other hand, the rare morph may suffer from pollen limitation due to their lower discovery and high inbreeding by pollinators, they seem to ensure reproductive success by producing higher quality nectar along with a higher self-compatibility rate. Since the higher quality nectar resulted in repeat visits by the pollinators most of which may be self-pollen (Harder *et al*., 2004; M. Eckhart *et al*., 2006; Ishii & Harder, 2006), presence of self-compatibility ensures that there is some seed-set, despite higher selfing. Thus, the plant’s reproductive fitness is correlated with morph abundance via both ecological and physiological paths suggesting a complex population-scale dynamics that may be responsible for the observed overall trimodal distribution of the morphs.

Population shifts in morph frequencies are common and have been observed in annual plants. However, across the last 15 years of our studies at the Kaas plateau (unpublished data, personal observations by authors) we did not record transitions in morph ratios or elimination of the rare morphs. This emphasizes the critical role vegetative propagation can play in maintaining rare morphs within a population. The importance of vegetative propagation has also been noted in other gingers (Záveská et al. 2012; Skopalíková et al., 2023; Yadav, 2025) and has been predicted to reduce the intensity of selection otherwise expected in sexually propagated plants (Jesson and Barrett 2002; Barrett and Harder 2006). Therefore, to understand factors shaping population structures of rhizomatic plants, genetic contributions from both sexual and vegetative strategies need to be considered in future studies.

Our study describes morph variability, specifically in reproductive outcomes, and our ongoing genetic studies are now focused on population genetic structure of the population as well as in identifying morph-level genotypic differences. Genetic analyses of population structure can help us partition the contributions of sexual versus vegetative reproduction and will hopefully aid in distinguishing relic genotypes from recent selection products (Eckert et al., 1999; Barrett, 2015). We aim to use *C. caulina* as a model species for long-term demographic studies focused on understanding polymorphism in perennial plants. By carrying out periodic plant and pollinator census, records of reproductive fitness of the morphs, and climate factors we hope to identify the transient or stable equilibria of the morph frequencies within and across populations of these endemic taxa from the lateritic plateaus.

## Conclusions

Our findings suggest a pluralistic view of floral polymorphism in perennial plants: pollinator visitation is a necessary but not sufficient predictor of morph fitness, because nectar economics, physiological compatibilities, mating differences, and demography jointly shape reproductive success. By foregrounding a multi-trait, network perspective we provide an empirical foundation for future examinations (paternity analysis, population genetic studies, and manipulative experiments) that will be required to separate contemporary selective forces from the legacies of the past. More broadly, studies of intraspecific polymorphism in long-lived plants will benefit from routinely measuring both pre- and post-pollination traits and from explicitly accounting for vegetative reproduction; only then can we robustly assess when and how color polymorphisms are maintained in nature.

## Supplementary Information

**Fig. S1** Plant habit, habitat, and key vegetative features of *Curcuma caulina*.

**Fig. S2** Variation in bract color, floral reflectance, and morph frequency of *C. caulina*.

**Fig. S3** Hawkmoth pollinators of *C. caulina*.

**Fig. S4** Circular histogram and circular statistics values of pollinator visitation rates.

**Table S1** List of floral and vegetative morphological characters used for the nMDS analysis.

**Table S2** Within-group comparisons for nectar energy, volume, and concentration.

**Table S3** Within-group comparisons for peak pollinator visitation rates.

**Table S4** Results of the chi-square test for fruiting success rates of hand-pollination.

**Table S5** Mean and standard error values for seed count for cross- and self-compatibility treatments.

**Table S6** Between groups comparisons for seed counts for hand pollination treatments.

## Author contributions

SS and VG conceived and designed the study. SS conducted the experiments and collected the field data. SS performed the statistical analyses with inputs from VG. SS drafted the manuscript; VG provided conceptual advice and edited the manuscript. All authors gave final approval for publication.

## Data availability

The entire dataset and analysis codes used in this study will be made accessible from Dryad or similar online repository after publication of the paper.

## Acknowledgements

Authors thank all funding agencies for funds to both authors and IISER Bhopal for infrastructure. We also thank PCCF, Maharashtra Forest Department, DFO of Satara Division (Forest Department), and the Joint Forest Management Committee (JFMC), Kaas, Satara for permitting our fieldwork. We thank Mr. Dilip Shrotri and Azad G for assistance during night work, Amal, Rahul, Viraj, Asawari, Najla, and Sooraj for their assistance during various data collections, including morphometry and Susnata and Sukhraj for assistance during physiological compatibility experiments during the fieldwork. Finally, we thank all fellow TrEE lab members for their valuable advice, discussions, and support.

