## Supplementary material for "A tale of three morphs I: Variations in nectar, reproductive compatibility, and abundance explain reproductive success of the morphs in a polymorphic ginger from Western Ghats, India"

### Annals of Botany: Supporting Information

Article acceptance date:

**Brief legends and index:**

**Fig. S1** Plant habit, habitat, and key vegetative features of *Curcuma caulina*.	2

**Fig. S2** Variation in bract color, floral reflectance, and morph frequency of *C. caulina*.	3

**Fig. S3** Hawkmoth pollinators of *C. caulina*.	4

**Fig. S4** Circular histogram and circular statistics values of pollinator visitation rates.	5

**Table S1** List of floral and vegetative morphological characters used for the nMDS analysis..	6

**Table S2** Within-group comparisons for nectar energy, volume, and concentration.	8

**Table S3** Within-group comparisons for peak pollinator visitation rates.	9

**Table S4** Results of the chi-square test for fruiting success rates of hand-pollination.	9

**Table S5** Mean and standard error values for seed count for cross- and self-compatibility treatments.	10

**Table S6** Between groups comparisons for seed counts for hand pollination treatments.	11

#### Fig. S1 (a) Mass flowering of *Curcuma caulina* with sympatric presence of bract colour morphs, in its native habitat at the study population, Kaas plateau (KAS). Inset shows the relative geographical location of KAS on the map of peninsular India. (b) Plant habit with key vegetative morphological characters.

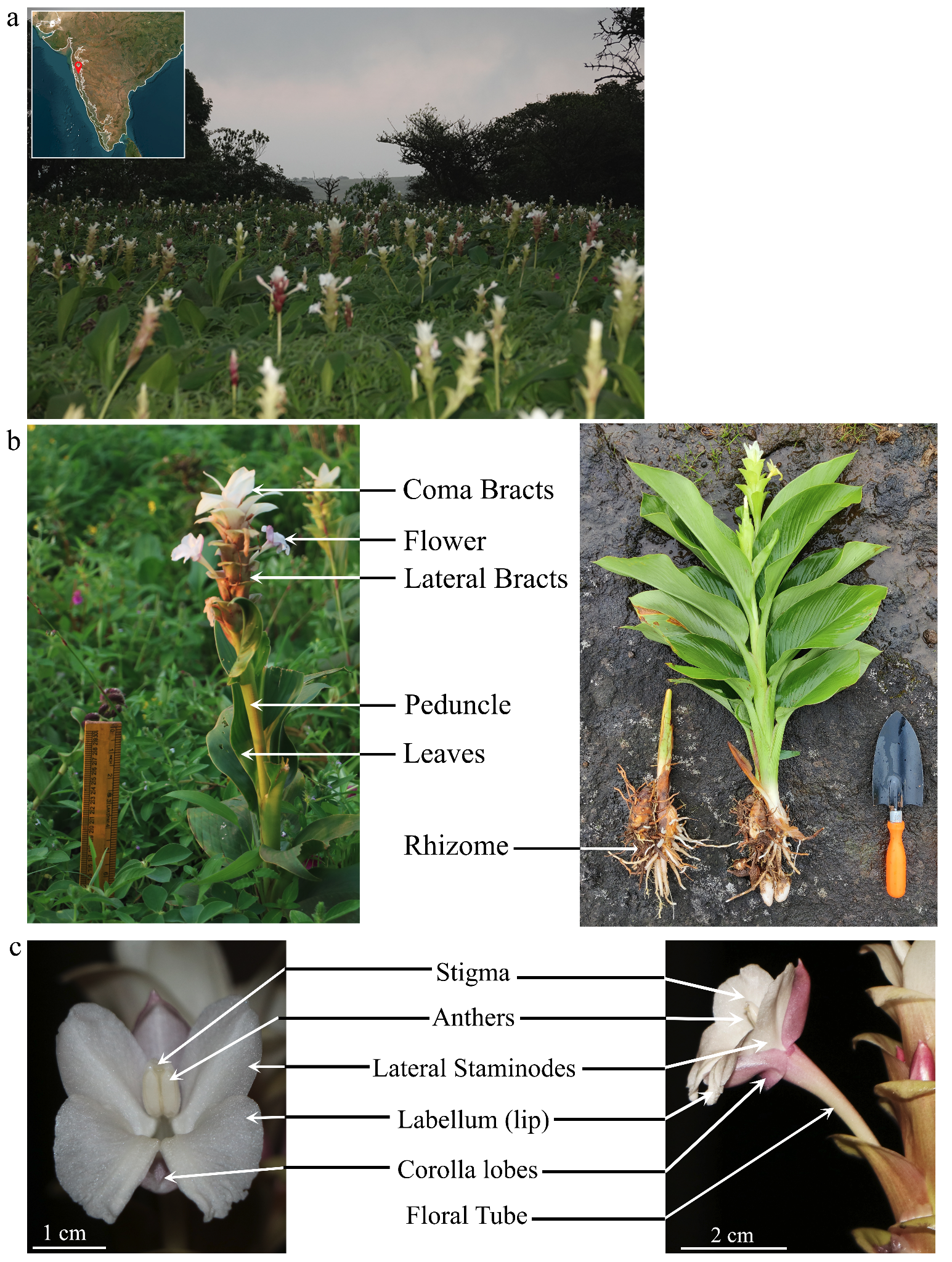

#### Fig. S2 (a) Sympatric bract color variation in *C. caulina*. Each inflorescence represents a different individual within the Kaas population. Note that flowers were consistently white in color, irrespective of the morph or variant type. We identified six color variants based on the percentage spread of red color on the first (or basal) lateral bract of the inflorescence and (b) the reflectance spectra of the basal lateral bracts for the three most abundant color variants (GR, GW, and RW). (c) Number of individuals of bract color variants of *C. caulina* present in ten transects (10 x 4 m) placed randomly across the study population. (d) Results of nMDS analysis for floral color morphs of *C. caulina* using 15 vegetative and 18 floral morphological characters. The analysis demonstrated lower stress values in a three-dimensional configuration (stress = 0.06) than in a two-dimensional configuration (stress = 0.11). The results suggest that bract color varies independently of the other morphological floral traits.

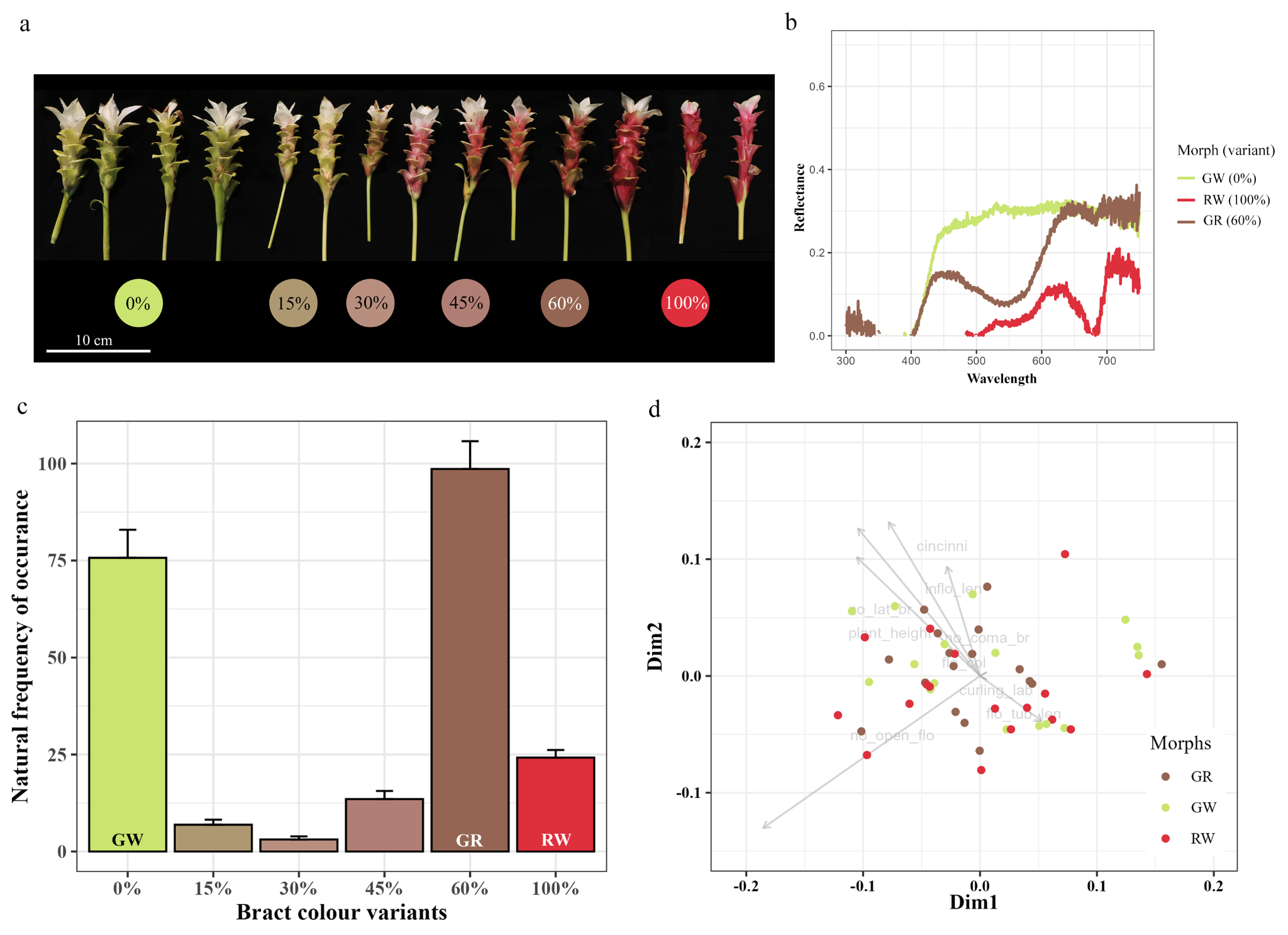

#### Fig. S3 Pollinators of *C. caulina*; a. Hawkmoth *Agrius convolvuli* visiting GR morph; b. Hawkmoth *Hippotion rafflesii* visiting RW morph, and c. pollen deposition site is the proboscis of the pollinator (shown by white arrowhead). These are the positions in which pollinators usually forage for nectar.

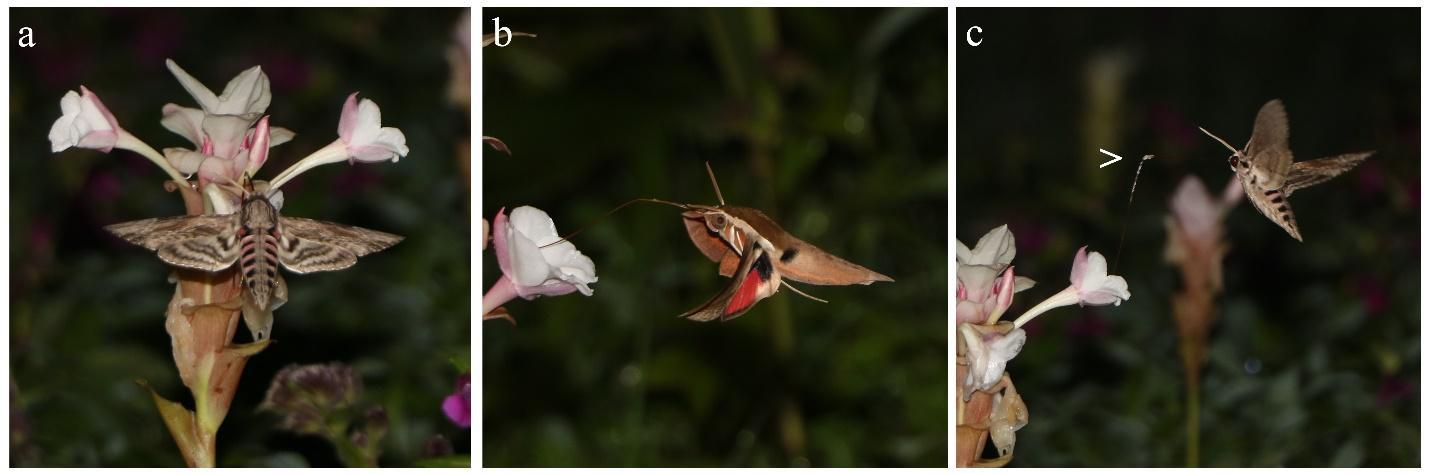

#### Fig. S4 Circular histogram of mean number of pollinator visits in the wild received by each morph. The 24 hours of a day are represented on the circular scale by angles with intervals of 15º. The direction of the vector (r) represents mean time and the length of this mean vector relative to the stippled line is a measure of concentration (K) of the data. The table below shows first order statistical values, results of Rayleigh test, and results of within-group comparisons (Mardia-Watson-Wheeler Test) for natural pollinator visitation rates for the three floral color morphs. The significance level was set at 0.05, with significant *p*-values indicated by an asterisk (*).

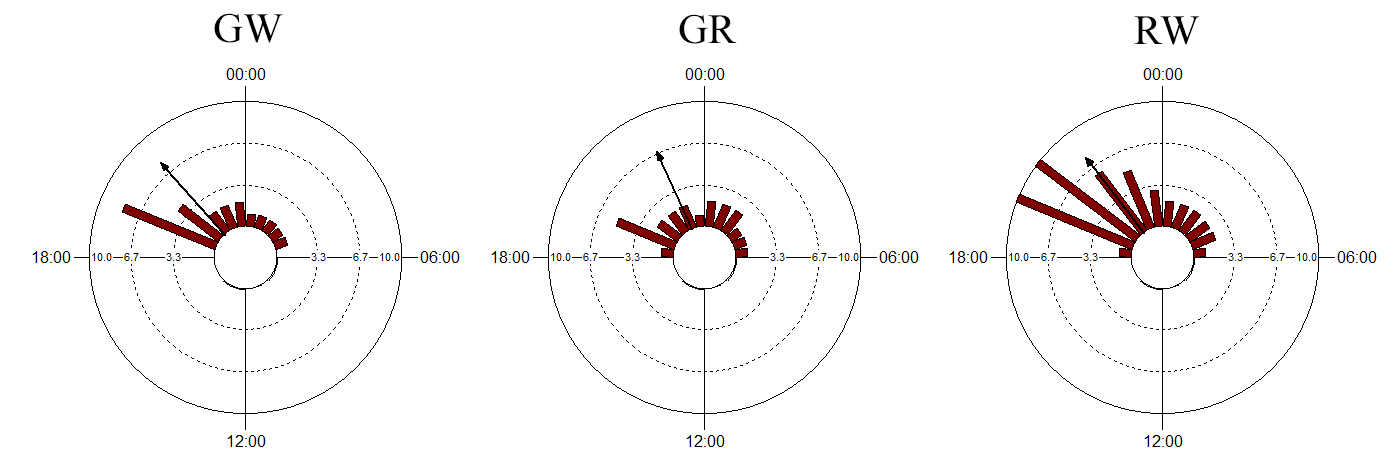

| **Morph** | **GW** | **GR** | **RW** |  | **GW** | **GR** | **RW** |
| --- | --- | --- | --- | --- | --- | --- | --- |
| Mean Vector (µ) | 21:07 (316.834°) | 22:00 (330.036°) | 21:21 (320.378°) | Rayleigh Test (Z) | 56.546 | 66.534 | 108.077 |
| Length of Mean Vector (r) | 0.776 | 0.718 | 0.786 | Rayleigh Test (p) | < 0.0001* | < 0.0001* | < 0.0001* |
| Median | 20:00 (300°) | 22:00 (330°) | 21:00 (315°) | **Mardia-Watson-Wheeler Test** | | | |
| Concentration | 2.594 | 2.124 | 2.7 |  | **W** | **p-value** | **Number of observations** |
| Circular Variance | 0.224 | 0.282 | 0.214 |  | **3.563** | **0.673** | **398** |
| Circular Standard Deviation | 02:43 (40.847°) | 03:06 (46.621°) | 02:39 (39.776°) |  | **GR** | **GW** | **RW** |
| Standard Error of Mean | 00:16 (4.166°) | 00:16 (4.084°) | 00:11 (2.973°) | **GR** | ----- | 0.122 | 0.227 |
| 95% Confidence Interval (-/+) for µ | 20:34 (308.667°) | 21:28 (322.029°) | 20:58 (314.55°) | **GW** | 4.47 | ----- | 0.313 |
|  | 21:40 (325.001°) | 22:32 (338.044°) | 21:44 (326.207°) | **RW** | 0.621 | 3.321 | ----- |

#### Table S1 List of floral and vegetative morphological characters used for the nMDS analysis. From the 33 morphometric characters, 22 characters were converted to ratios in order to avoid pseudoreplication in characters. To avoid multiple correlated variables (denoted by * in the first column), we took ratios of paired variables. Characters which are used in actual analysis are given in the third column and abbreviations for these characters are given in the fourth column.

| **Sr No.** | **Characters** | **Used in the analysis as** | **Abbreviation used in the analysis** |
| --- | --- | --- | --- |
| 1 | Plant height | Plant height | plant_height |
| 2 | Number of leaves | Number of leaves | no_leaves |
| 3* | Lamina length | Ratio (lamina length/lamina width) | lam_ratio |
| 4* | Lamina width |  |  |
| 5* | Inflorescence length | Ratio (inflorescence length/ inflorescence width) | inflo_ratio |
| 6* | Inflorescence width |  |  |
| 7 | Number of open flowers | Number of open flowers | no_open_flo |
| 8* | Peduncle length | Ratio (peduncle length/ peduncle width) | ped_ratio |
| 9* | Peduncle width |  |  |
| 10 | Colour at base of the pseudostem | Colour at base of the pseudostem | col_at_base |
| 11 | Number of coma bracts | Number of coma bracts | no_coma_br |
| 12 | Number of lateral bracts | Number of lateral bracts | no_lat_br |
| 13 | Length of the lateral bracts | Length of the lateral bracts | lat_br_len |
| 14 | Width of the lateral bract | Width of the lateral bract | lat_br_wid |
| 15 | Number of cincinni | Number of cincinni | cincinni |
| 16* | Bracteole length | Ratio (bracteole length/ bracteole width) | brctol_ratio |
| 17* | Bracteole width |  |  |
| 18* | Calyx length | Ratio (calyx length/ calyx width) | cal_ratio |
| 19* | Calyx width |  |  |
| 20* | Corolla tube length | Ratio (corolla tube length/ corolla tube width) | cor_tub_ratio |
| 21* | Corolla tube width |  |  |
| 22* | Corolla lobe length | Ratio (corolla lobe length/ corolla lobe width) | cor_lob_ratio |
| 23* | Corolla lobe width |  |  |
| 24* | Lateral staminode length | Ratio (lateral staminode length/ lateral staminode width) | lat_st_ratio |
| 25* | Lateral staminode width |  |  |
| 26* | Labellum length | Ratio (labellum length/ labellum width) | lab_ratio |
| 27* | Labellum width |  |  |
| 28 | Depth of the notch | Depth of the notch | notch_dep |
| 29* | Anther length | Ratio (anther length/ anther width) | anth_ratio |
| 30* | Anther width |  |  |
| 31 | Style length | Style length | style_len |
| 32* | Nectaries length | Ratio (nectaries length/ nectaries width) | nect_ratio |
| 33* | Nectaries width |  |  |

#### Table S2 Results of within-group comparisons for (a) nectar energy (cal), (b) nectar volume and (c) nectar concentration across three morphs of *C. caulina*. Significance was set at 0.05, with significant *p*-values indicated by an asterisk (*).

**a)**

| **Morph** | **Nectar volume (μl)** | |  |  |  |  |  |  |
| --- | --- | --- | --- | --- | --- | --- | --- | --- |
|  | **Mean** | **SEM** | **Kruskal-wallis chi squared** | **df** | ***p*-value** | **Comparison between groups (Dunn’s test)** | | |
| GR | 16.7368 | 1.473 | 33.487 | 2 | <0.0001* |  | **GR** | **GW** |
| GW | 18.0489 | 2.227 |  |  |  | **GW** | 0.3178 |  |
| RW | 31.3075 | 2.253 |  |  |  | **RW** | <0.0001* | <0.0001* |

**b)**

| **Morph** | **Nectar volume (μl)** | |  |  |  |  |  |  |
| --- | --- | --- | --- | --- | --- | --- | --- | --- |
|  | **Mean** | **SEM** | **Kruskal-wallis chi squared** | **df** | ***p*-value** | **Comparison between groups (Dunn’s test)** | | |
| GR | 11.8296 | 0.866 | 32.619 | 2 | <0.0001* |  | **GR** | **GW** |
| GW | 13.2484 | 1.375 |  |  |  | **GW** | 0.1936 |  |
| RW | 21.0043 | 1.38 |  |  |  | **RW** | <0.0001* | <0.0001* |

**c)**

| **Morph** | **Nectar concentration (% sucrose)** | |  |  |  |  |  |  |
| --- | --- | --- | --- | --- | --- | --- | --- | --- |
|  | **Mean** | **SEM** | **Kruskal-wallis chi squared** | **df** | ***p*-value** | **Comparison between groups (Dunn’s test)** | | |
| GR | 30.7053 | 0.654 | 10.13 | 2 | 0.006315* |  | **GR** | **GW** |
| GW | 28.6438 | 0.967 |  |  |  | **GW** | 0.0498* |  |
| RW | 32.5056 | 0.504 |  |  |  | **RW** | 0.0521 | 0.0007* |

#### Table S3 Results of within-group comparisons for peak pollinator visitation rates (# visits per flower per hour) for all three morphs of *C. caulina*. Significance was set at 0.05, with significant *p*-values indicated by an asterisk (*).

| **Kruskal-wallis chi squared** | **df** | ***p*-value** | **Comparison between groups (Dunn’s test)** | | |
| --- | --- | --- | --- | --- | --- |
| 6.52 | 2 | 0.03839* |  | **GR** | **GW** |
|  |  |  | **GW** | 0.3377 |  |
|  |  |  | **RW** | 0.0066* | 0.0302* |

#### Table S4 Results of chi-square test for success rates of self- and cross-compatibility treatments conducted for all three morphs. Significance was set at 0.05, with significant *p*-values indicated by an asterisk (*).

| **Treatment** | **Effect** | **Chi-square value** | **df** | ***p*-value** |
| --- | --- | --- | --- | --- |
| **All three** | **Treatments** | 29.278 | 1 | < 0.0001* |
| **Self-pollination** | **Female morph** | 2.8681 | 2 | 0.2383 |
| **Intra-morph cross-pollination** | **Male/ female morph** | 5.5019 | 2 | 0.06387 |
| **Inter-morph cross-pollination** | **Pair a: GW and GR morphs** | 1.5771 | 1 | 0.2092 |
|  | **Pair b: GW and RW morphs** | 2.2711 | 1 | 0.1318 |
|  | **Pair c: GR and RW morphs** | 0.22996 | 1 | 0.6316 |

#### Table S5 Mean and standard error (s.e.) values for seed-count for cross- and self-compatibility treatments conducted for all three morphs.

| **Male** | **Female** | **Mean** | **sem** |
| --- | --- | --- | --- |
| **Self-pollination treatments** | | | |
| Self | GR | 0 | 0 |
| Self | GW | 14.5 | 6.5 |
| Self | RW | 13.33333333 | 4.371625683 |
| **Intra-morph cross-pollination treatments** | | | |
| GR | GR | 20.71428571 | 2.825372817 |
| GW | GW | 10.16666667 | 2.713136766 |
| RW | RW | 15.7 | 2.712317582 |
| **Inter-morph cross-pollination treatments** | | | |
| GR | GW | 20.84615385 | 2.556655853 |
| GR | RW | 20.42857143 | 3.294541416 |
| GW | GR | 15 | 2.872281323 |
| GW | RW | 12.125 | 2.552572232 |
| RW | GR | 21.5 | 3.12161905 |
| RW | GW | 17.8125 | 2.770708017 |

#### Table S6 Results of between groups comparisons for seed-count for self- and cross-compatibility treatments for all three morphs. Significance was set at 0.05, with significant *p*-values indicated by an asterisk (*).

| **A. For self-pollination treatments** | | | | |
| --- | --- | --- | --- | --- |
| **Kruskal-Willis chi-squared = 5.3534, df = 2, *p*-value = 0.06879** | | | | |
| **Female morphs** |  | **GR** | **GW** |  |
|  | **GW** | 0.0419* |  |  |
|  | **RW** | 0.0158* | 0.4238 |  |
| **B. For intra-morph cross-pollination treatments** | | | | |
| **Kruskal-Willis chi-squared = 5.2499, df = 2, *p*-value = 0.07244** | | | | |
|  |  | **GR x GR** | **GW x GW** |  |
|  | **GW x GW** | 0.0115* |  |  |
|  | **RW x RW** | 0.1419 | 0.0987 |  |
| **C. For inter-morph cross-pollination treatments** | | | | |
| **Pairs** | | **a** | **b** | **c** |
| **Kruskal-Willis chi-squared** | | **1.5177** | **1.3614** | **0.03389** |
| **df** | | **1** | **1** | **1** |
| ***p*-value** | | **GR x GW** | **RW x GW** | **RW x GR** |
|  | **GW x GR** | 0.218 |  |  |
|  | **GW x RW** |  | 0.2433 |  |
|  | **GR x RW** |  |  | 0.8539 |
